# ETS1 Orchestrates a Hybrid EMT Program Driving in vivo Metastasis and Immune Evasion

**DOI:** 10.1101/2025.07.17.665404

**Authors:** Benjamin Ziman, Talia A. Wenger, Chehyun Nam, Qian Yang, Daniel Arnaudov, Megha Sheth, Zhixuan Jing, Yuhao Pan, Joe Vargas, Yong Teng, Uttam K. Sinha, Young Min Park, De-Chen Lin

## Abstract

Transcriptional Intratumoral heterogeneity (ITH) is a hallmark of aggressive cancers, yet how transcriptional ITH programs drive tumor metastasis and immune evasion in upper aerodigestive squamous cell carcinoma (UASCC) remains unclear. Through single-cell RNA sequencing analysis of UASCC cells and patient tumors, we uncovered a hybrid epithelial mesenchymal transition (hEMT) ITH program linked to metastatic dissemination. The transcription factor ETS1 was identified as a master regulator of the hEMT program, directly activating pro-metastatic genes and promoting distant spread in vivo. Unexpectedly, ETS1 also orchestrated an immune-cold tumor microenvironment by transcriptionally activating both *STAT1* and *PD-L1 (CD274)* genes, suppressing T lymphocyte infiltration, and elevating immune checkpoint molecules. Clinically, ETS1-high tumors strongly correlated with poor survival and resistance to immune checkpoint blockade across multiple cohorts. Leveraging drug screens, we discovered that ETS1-high cancers are vulnerable to HSP90 inhibitors (e.g., Alvespimycin), which suppress ETS1 by disrupting HIF1A-mediated transcriptional activation. Together, our work reveals ETS1 as a dual driver of tumor distal metastasis and immune evasion in UASCC, while nominating HSP90 inhibition as a tailored treatment strategy for ETS1-driven tumors. These findings provide a roadmap for targeting aggressive ITH subsets and overcoming immunotherapy resistance.

## Introduction

Upper aerodigestive squamous cell carcinoma (UASCC) encompasses a group of aggressive malignancies originating from the epithelial lining of the oral cavity, pharynx, larynx, and esophagus. Over one million new cases of UASCC are diagnosed worldwide every year, with head and neck squamous cell carcinoma (HNSCC) and esophageal squamous cell carcinoma (ESCC) being the most common types^1^. Despite advances in surgical and therapeutic interventions, patient outcomes remain poor, particularly for those who develop distal metastases. Tumor metastasis to distant organs such as lung and liver represents the most significant clinical challenge in UASCC, as it is the leading cause of treatment failure and cancer-related mortality^2^. To date, there are no effective approaches to prevent metastatic progression, underscoring the urgent need to unravel the underlying molecular mechanisms driving this phenomenon.

A major obstacle in addressing cancer metastasis lies in the extensive transcriptional intratumoral heterogeneity (ITH) of UASCC. Tumors exhibit diverse cellular processes and biological programs that are linked to critical cancer hallmarks, including drug resistance, immune evasion, and metastatic potential^3^. These ITH-driven cellular programs enable tumor cells to adapt to selective pressures from therapy and from the microenvironment through the creation of subpopulations with specialized functions that contribute to disease progression and treatment failure. Characterizing these transcriptional ITH programs at a high resolution is essential for developing novel therapeutic strategies.

Advances in single-cell RNA sequencing (scRNA-seq) have revolutionized our ability to dissect ITH programs at an unprecedented scale^4–8^. By capturing the transcriptional profiles of individual cells within the tumor, scRNA-seq enables the identification of distinct cellular subpopulations and their associated gene expression programs. Computational approaches, such as non-negative matrix factorization (NMF), further enhance these efforts by delineating transcriptional gene modules that define key functional states, such as those driving cellular stress, senescence, and epithelial-mesenchymal transition (EMT)^4–9^. Of note, a partial- or hybrid-EMT state, characterized by the coexistence of epithelial and mesenchymal markers, as well as a lack of canonical EMT transcription factors, has been discovered in UASCC^4–6,9^.

In addition to intrinsic tumor programs, the interaction between cancer cells and the immune microenvironment profoundly influences UASCC progression. Immune checkpoint blockade (ICB) therapies have emerged as a promising strategy to reinvigorate anti-tumor immunity in cancer patients. However, the efficacy of ICB in UASCC is often limited^10,11^, likely due to the immunosuppressive milieu driven by tumor heterogeneity. While not fully understood, distal metastasis appears to be intricately linked to tumor immune evasion. Indeed, emerging evidence suggests that metastatic cells may evade immune surveillance by modulating immune checkpoint pathways, suppressing antigen presentation, or creating an immunosuppressive microenvironment in distal organs^12–14^. However, the specific molecular and cellular mechanisms enabling this immune escape during metastasis are not fully elucidated in UASCC. Understanding these processes is crucial for improving therapeutic strategies, including the development of immune-based interventions to target metastatic disease.

To address these critical gaps, we performed integrative single-cell and functional analyses aimed at uncovering the key ITH programs, transcriptional regulators and vulnerabilities driving UASCC metastasis and progression. Our study focused on defining the molecular programs that coordinate metastasis and immune evasion, with the goal of identifying actionable targets for intervention. Here, we present our findings that position ETS1 as a central orchestrator of these malignant processes and reveal novel therapeutic opportunities for ETS1-high tumors.

## Results

### Identification of an intratumoral heterogeneity (ITH) transcriptional program associated with metastatic potential in UASCC

To identify ITH transcriptional programs associated with distal metastasis in UASCC, we began by leveraging a published in vivo study, Metmap, that profiled the metastatic potential of over 500 cancer cell lines^15^. Among these, 16 UASCC cell lines with matched scRNA-seq data^5^ were selected for further analysis. Recognizing the emerging role of gene modules— coordinately expressed sets of genes—as defining features of cellular states, we applied non-negative matrix factorization (NMF) to uncover transcriptionally distinct subpopulations within these cell lines (see Methods). NMF revealed seven clusters of recurrent gene modules, representing ITH transcriptional programs across individual cell lines (**Figures 1A-B**). These consensus programs were defined by sets of genes consistently upregulated in specific subpopulations, indicating heterogeneously expressed transcriptional profiles.

**Figure 1.**
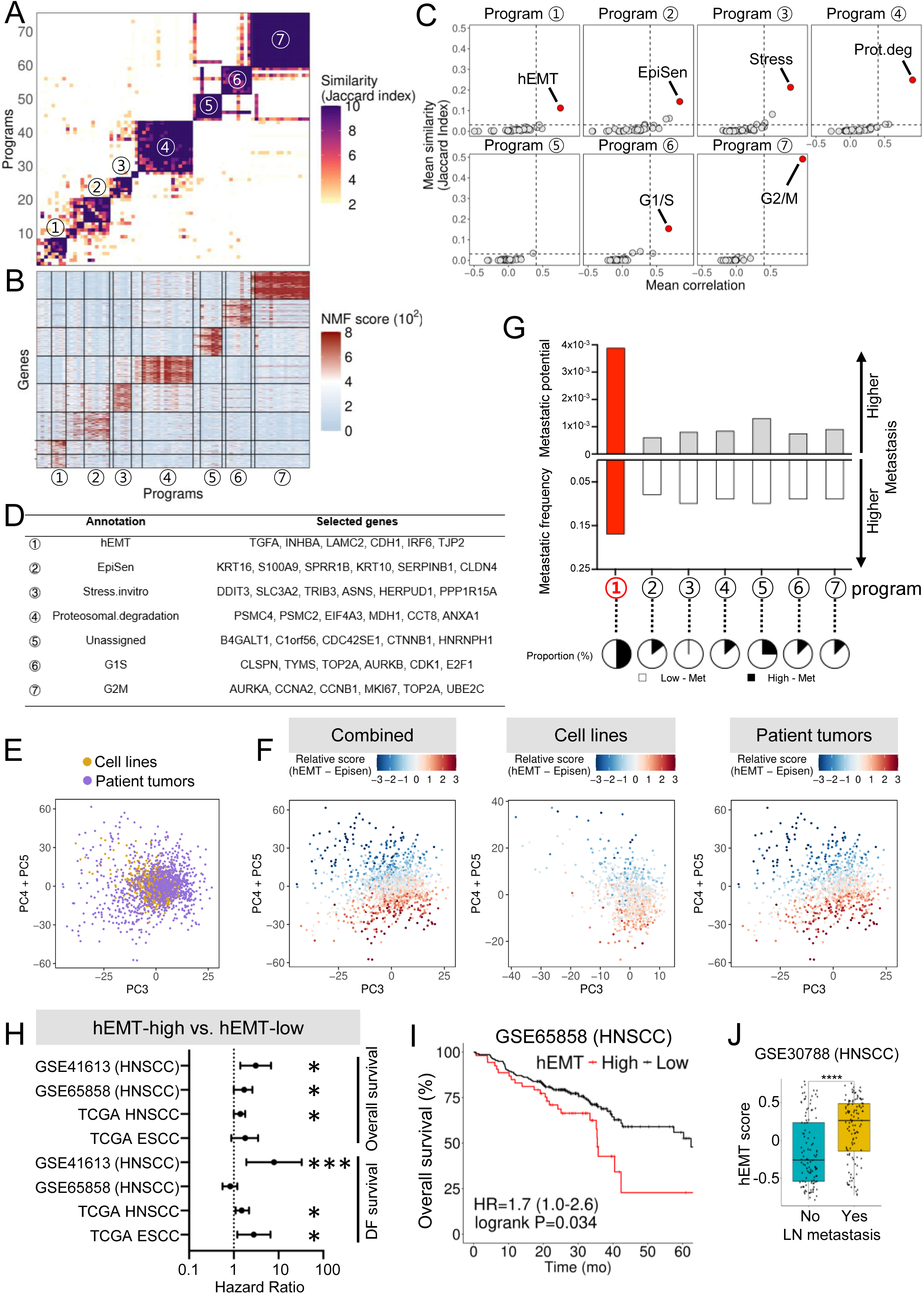
Identification of an intratumoral heterogeneity program associated with metastatic potential in UASCC. (A) Heatmap showing unsupervised clustering of 50 NMF programs based on their top genes, with similarity measured by Jaccard indices. Seven consensus ITH program clusters are numbered and highlighted. (B) Heatmap displaying the expression levels of program genes aligned with the clusters shown in panel (A). (C) Dot plots illustrating the Jaccard index (y-axis) and mean correlation (x-axis) between ITH programs and published meta-programs^5^. (D) Representative gene lists of each ITH program from panel (B). (E) PCA plots of combined cell lines and primary HNSCC tumor cells. (F) Cells are color-coded by the relative score for the hEMT minus EpiSen program. (G) Column plot showing the mean values of metastatic potential and penetrance for cell lines enriched in each ITH program. (H) Line plot showing the hazard ratio and p-values for the hEMT gene signature in the indicated UASCC clinical cohorts, using the singscore method^40^. (I) Representative Kaplan-Meier (KM) plots from panel (H). (J) Box plot showing the expression of the hEMT gene signature in HNSCC tumors with or without lymph node metastasis.

To understand the functional roles of the seven ITH programs identified, we performed enrichment analysis using well-curated gene programs derived from a similar NMF-based approach on large-scale scRNA-seq data from pan-cancer tumor samples^6^. This analysis revealed distinct biological processes associated with each program (**Figures 1C-D**). Specifically, Program-2 was annotated as an epithelial senescence (EpiSen) program, while Programs 3 and 4 corresponded to stress response and proteasomal degradation, respectively. Programs 6 and 7 were associated with cell cycle progression, specifically G1/S and G2/M phases, respectively. While Program-1 was enriched with genes from a pan-cancer EMT program, it also notably included epithelial differentiation genes such as CDH1, IRF6, JUP, KLF6, and TJP2 (**Figure 1D, Supplementary Table 1**). The coexistence of EMT-associated and epithelial-specific features strongly resembled the partial or hybrid EMT (hEMT) programs previously identified in HNSCC^4–6^. As such, Program-1 was annotated as an hEMT program.

To validate that the ITH programs identified in UASCC cell line models reflect those expressed in patient tumors, we conducted a combined analysis of cell line scRNA-seq data with in vivo scRNA-seq data from HNSCC fresh tumors^4^. PCA plots revealed that in vitro tumor cells and in vivo HNSCC primary cells shared overlapping transcriptional space (**Figure 1E**), indicating high comparability between these models. We then plotted relative scores for the hEMT and EpiSen programs at the single-cell level, focusing on these programs since EMT and EpiSen represent two polarized states of intratumoral malignant cells^5^. Indeed, cells exhibiting hEMT or EpiSen signatures were distinctly separated, occupying opposite regions in a comparable pattern between cell lines and in vivo tumors (**Figure 1F**). These findings confirm that the ITH programs identified in UASCC cell lines robustly recapitulate the cellular states and transcriptional heterogeneity of UASCC patient tumors, reinforcing the validity of our approach for studying intratumoral dynamics.

We next explored whether any of the seven ITH programs were associated with the distant metastatic capability of UASCC cell lines. Specifically, we examined the mean metastatic potential (quantified as the percentage of metastatic cells detected in vivo^15^) and the mean metastatic penetrance (defined by the percentage of animals developing metastasis^15^) of the cell lines classified under each ITH program. Notably, cell lines assigned to the hEMT program exhibited the highest levels of both metastatic potential and penetrance, **Figure 1G**). This finding suggests that the hEMT program may play a crucial role in promoting distant metastatic behavior in UASCC. Congruently, using the hEMT program gene signature, we found strong correlations with poor patient survival in both HNSCC and ESCC across multiple independent clinical cohorts (**Figures 1H-I**). Additionally, the hEMT signature was associated with tumor lymph node metastasis in independent HNSCC datasets (**Figure 1J**).

### ETS1 is a key regulator of the hEMT gene program and is associated with UASCC metastasis

To identify potential upstream regulators of hEMT program genes, we performed SCENIC^16^ regulon analysis, a robust method for linking regulatory factors to gene expression modules by integrating single-cell transcriptomic data and motif sequence enrichment. Analysis of three highly metastatic UASCC cell lines associated with the hEMT program revealed two shared transcription factors, ETS1 and ELK3 (**Supplementary Figure 1A**). ETS1 was prioritized for downstream analysis due to its significant overexpression in UASCC tumors compared to nonmalignant adjacent tissues (**Supplementary Figure 1B**).

To validate the SCENIC regulon data derived from UASCC cell lines, we first examined epithelial-specific ETS1 expression in a combined scRNA-seq HNSCC cohort, comprising both public datasets^4,17–20^ and internal patient samples (total n=114). This independent analysis confirmed a significant correlation between ETS1 expression and the hEMT program score across the HNSCC scRNA-seq dataset (**Figure 2A**). To further substantiate this finding, we analyzed three independent bulk RNA-Seq datasets: UASCC cell lines from Cancer Cell Line Encyclopedia (CCLE; n=99), The Cancer Genome Atlas (TCGA) HNSCC tumors (n=520), and HNSCC patient-derived organoids (n=21) reported by us^9^. Notably, ETS1 expression consistently demonstrated a strong correlation with the hEMT program score across all three cohorts (**Figures 2B-D**), strongly suggesting ETS1 as a potential functional regulator of the hEMT program.

**Figure 2.**
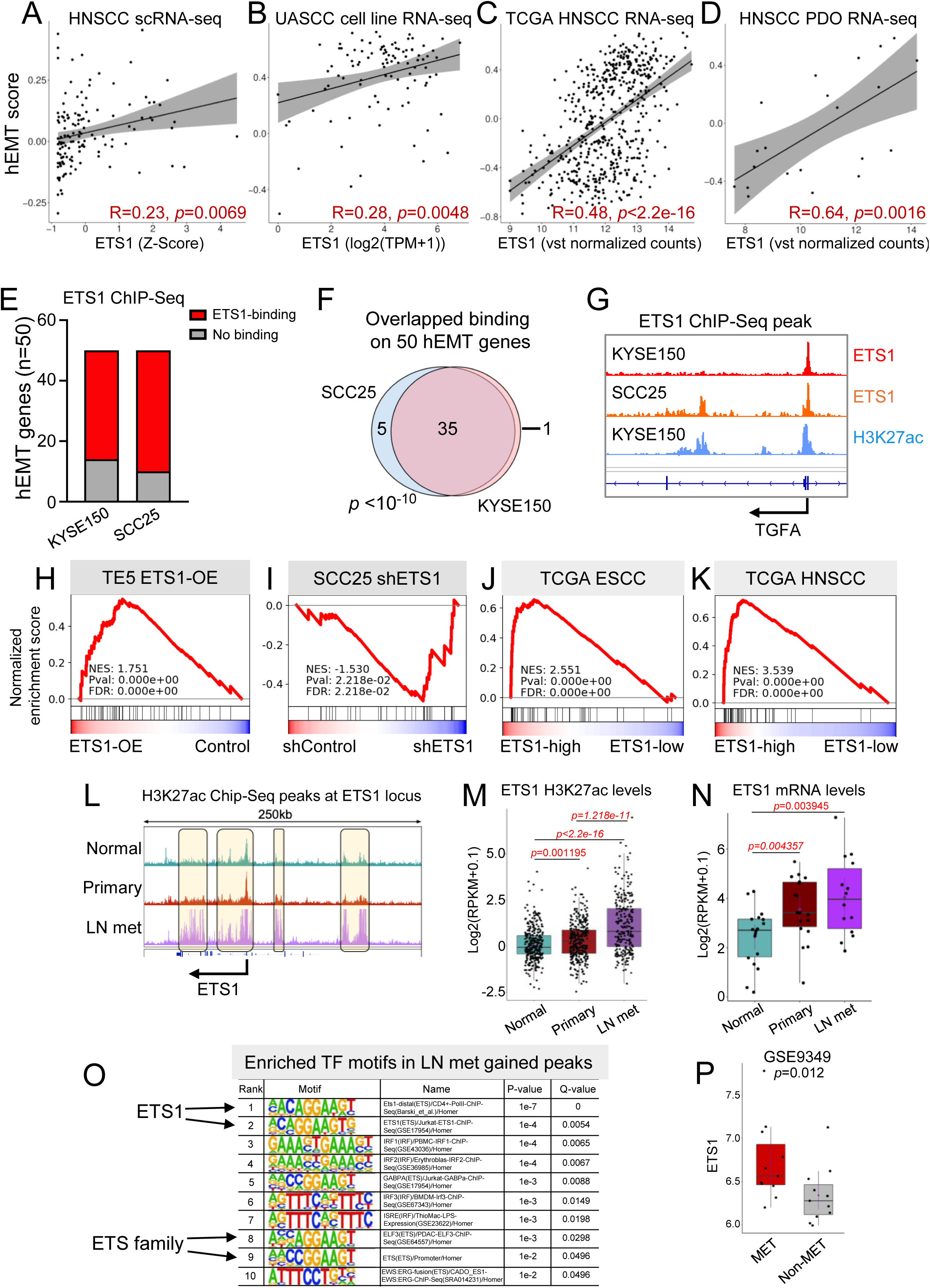
ETS1 is a key regulator of the hEMT gene program and is associated with UASCC metastasis. (A-D) Scatter plots showing correlation between ETS1 expression and the hEMT score in (A) human HNSCC scRNA-seq cohort, (B) UASCC cell lines, (C) TCGA HNSCC tumors, (D) HNSCC PDO samples. (E) Stacked plot illustrating ETS1 ChIP-Seq peaks on hEMT genes from two cell lines. (F) A Venn diagram showing shared ETS1 ChIP-Seq peaks between the KYSE150 and SCC25 cell lines, with representative peaks shown in (G). (H-K) GSEA line plots showing enrichment of the hEMT gene signature following ETS1 overexpression (H) or knockdown (I), or in ETS1-high tumors within TCGA ESCC (J) and HNSCC (K) cohorts. (L) An IGV plot showing H3K27ac ChIP-Seq profiles in nonmalignant esophageal tissues, primary ESCC, and lymph node (LN) metastases (PRJNA665151), with quantification shown in panel (M). (N) ETS1 mRNA expression in the same samples as panel (L) and in GSE9349 (P) dataset. (O) Motif sequence enrichment analysis of gained H3K27ac peaks in LN metastases from PRJNA665151.

We next performed ChIP-Seq using an ETS1 antibody in KYSE150, a highly metastatic UASCC line based on the MetMap^15^. This analysis revealed that 36 of the 50 hEMT genes contained ETS1-binding peaks, suggesting direct transcriptional regulation (**Figures 2E, G**). To corroborate these findings, we analyzed publicly available ETS1 ChIP-Seq data^21^ from SCC25, another metastatic UASCC line. Importantly, 40 of the 50 hEMT genes displayed ETS1 occupancy in SCC25, with striking overlap observed between these two datasets (**Figures 2F-G**).

To further verify ETS1’s role in regulating the hEMT program, we ectopically overexpressed ETS1 in the TE5 cell line, which has low metastatic capability (based on the MetMap data^15^, **Supplementary Figure 2A**) and low ETS1 expression (based on the CCLE data^22^, **Supplementary Figure 2B**). RNA-Seq analysis demonstrated significant upregulation of the hEMT gene-set upon ETS1-overexpression, as evidenced by the GSEA result (**Figure 2H**). Conversely, knockdown of ETS1 in the ETS1-high line SCC25 produced the opposite effect (**Figure 2I**), providing functional evidence of ETS1’s regulatory influence on the hEMT program in UASCC cells. We next analyzed RNA-Seq datasets from the TCGA ESCC and HNSCC cohorts after stratifying patient samples into ETS1-high and ETS1-low groups based on ETS1 expression levels. Consistently, the hEMT gene set was strongly enriched in the ETS1-high group (**Figure 2J-K**).

To explore the epigenetic basis of ETS1 activation during metastasis, we analyzed enhancer and promoter landscapes from an epigenomic dataset comparing nonmalignant esophageal tissues, primary ESCC, and LN metastases^23^. Notably, H3K27ac deposition at the ETS1 locus was substantially higher in LN metastases than in either nonmalignant samples or primary ESCC tumors, pointing to epigenetic activation of ETS1 during lymphatic metastasis (**Figures 2L-M**). This activation aligns with the highest ETS1 mRNA levels in LN metastases in two independent datasets (**Figures 2N-P**). To gain deeper insights into the regulatory landscape of metastatic progression, we performed differential H3K27ac peak analysis between primary ESCC and LN metastases, followed by transcription factor motif enrichment analysis (**Supplementary Figure 1C**). Notably, ETS1 motifs were the most significantly enriched within the gained peaks in LN metastases compared to primary tumors (**Figure 2O**), suggesting a pivotal role for ETS1 in epigenetic reprogramming associated with UASCC metastasis.

### ETS1 drives hEMT gene expression and promotes in vivo metastasis

To investigate the functional role of ETS1 in tumor metastasis, we selected two UASCC cell lines, TE5 and TT, with low ETS1 expression and relatively low metastatic potential for gain-of-function studies. (**Supplementary Figures 2A-B**). Conversely, for loss-of-function experiments, we used KYSE150, which exhibits high ETS1 expression and relatively high metastatic potential (**Supplementary Figures 2A-B**). We first validated the expression levels of hEMT-related genes in vitro. Using a randomly selected hEMT gene set comprising both canonical EMT genes (e.g., TGFA, LAMC2, LAMB3) and epidermal differentiation genes (e.g., CDH1, JUP, TJP2), we observed their increased expression upon ETS1 overexpression and reduced expression following ETS1 knockdown (**Figures 3A**).

**Figure 3.**
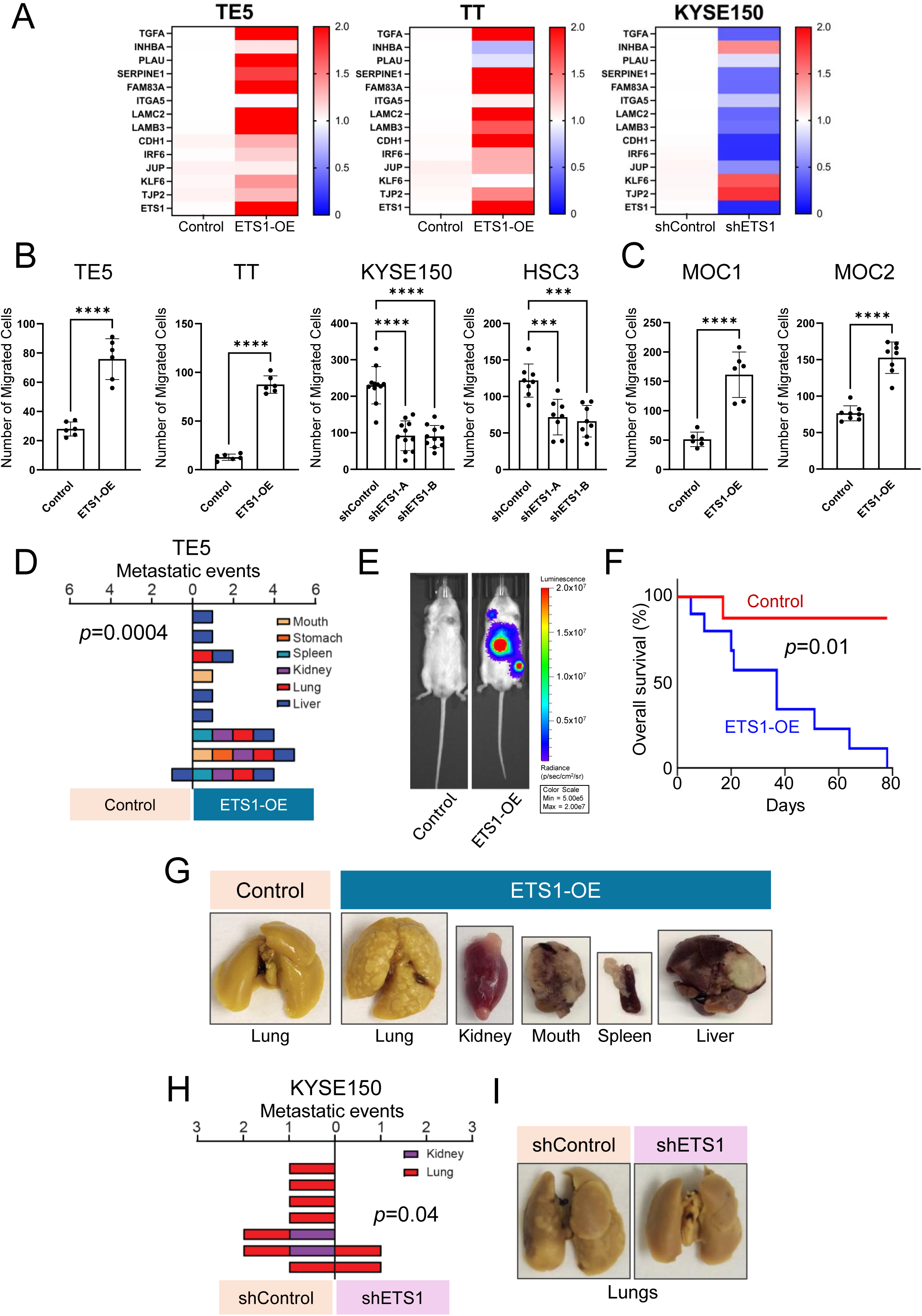
ETS1 is a regulator of hEMT gene expression and cell migration. (A) Heatmaps showing the relative expression of hEMT genes in UASCC cells. Data are shown as the mean of three experimental replicates. Expression is shown from high (red) to low (blue). Values expressing a fold change of 2 or greater are shown as the highest value (red). (B-C) Transwell migration assays of ETS1-overexpressing or ETS1-knockdown UASCC cells. Bar graphs show the number of migrated cells and represent the mean ± SD of three experimental replicates. ***P<0.001, ****P<0.0001; P-values were determined using an unpaired t-test or one-way ANOVA with multiple comparisons. (D) The metastatic events across various target organs in mice bearing stable control or ETS1-overexpressing TE5 cells. The number of metastatic events (grouped by location of metastasis) is shown for each mouse (horizontally) across the experimental (right) and control (left) groups. Target organs are visualized according to color; mouth (tan), stomach (orange), spleen (teal), kidney (purple), lung (red), and liver (blue). (E) Representative bioluminescence image depicting the spread of control and ETS1-overexpressing TE5 cells transfected with luciferase 21 days post-injection. (F) Kaplan-Meier plots showing mice bearing either control (red) or ETS1-ovexpressing (blue) cells. P-value was determined by the log-rank test. (G) Representative images of metastases to various organs of mice bearing control and ETS1-overexpressing TE5 cells. (H) The metastatic events across target organs in mice bearing control or stable ETS1-knockdown KYSE150 cells. (I) Representative images of the lungs of mice bearing control and stable ETS1-knockown KYSE150 cells.

Next, we assessed whether altering ETS1 levels impacts cell migration properties. Transwell migration assays revealed ETS1 overexpression significantly enhanced cell migration towards nutrient-rich areas, whereas ETS1 knockdown reduced migration (**Figure 3B**). These findings were corroborated in murine HNSCC cell lines MOC1 and MOC2, where increased ETS1 expression led to a higher number of migrated cells (**Figure 3C**). To further examine ETS1’s role in cell motility, we performed wound healing assays, confirming that ETS1 overexpression enhanced cell migration (**Supplementary Figure 2C**), while ETS1 knockdown diminished migration (**Supplementary Figure 2D**). These results were consistent in MOC1 and MOC2, where ETS1 overexpression also promoted migration (**Supplementary Figure 2E**). Collectively, these data suggest that ETS1 not only regulates the expression of the hEMT gene program but also drives phenotypic changes that enhance migratory capacity.

We next investigated the role of ETS1 in promoting cancer metastasis in vivo. We used the TE5 cell line to test whether ETS1 overexpression could enhance metastatic potential in cells that are otherwise weakly metastatic. To enable tumor tracking, we engineered luciferase-labeled cells. Either control or ETS1-overexpressing (ETS1-OE) TE5 cells were injected into the tail vein of immunodeficient mice. Strikingly, while only 11% of mice injected with control cells developed detectable metastases, 100% of mice injected with ETS1-overexpressing cells exhibited widespread metastatic disease (P = 0.0004; **Figures 3D-E**). In these mice, metastases involved many organs, including the lung, liver, kidney, stomach, spleen, etc. (**Figure 3D, G**). Moreover, ETS1 overexpression significantly reduced overall survival compared to the control group (**Figure 3F**).

In a complementary loss-of-function experiment, we assessed the impact of ETS1 knockdown in the highly metastatic KYSE150 cell line. It was observed that the majority of mice injected with control cells developed distal metastases (87.5%), whereas only 25% of mice injected with ETS1-knockdown cells exhibited metastatic disease (P = 0.04; **Figure 3H**). The lungs were the predominant site of metastasis, followed by the kidneys (**Figures 3H-I**). Together, these results demonstrate that ETS1 is a critical driver of metastasis in vivo, promoting both metastatic burden and disease severity across multiple organ sites.

### ETS1-high cancer vulnerability to HSP90 inhibition is mediated through HIF1A suppression

We next sought to explore the specific vulnerability of ETS1-high cancers by conducting an in silico screen based on the PRISM Primary Repurposing dataset^24^, in which 4,687 compounds were tested across 483 cancer cell lines with matched RNA-Seq data. By correlating ETS1 expression levels with compound efficacy, we identified top candidates— including HMGCR, HSP90, and Na⁺/K⁺-ATPase inhibitors—that demonstrated greater potency in killing ETS1-high cell lines (**Figure 4A**). To validate these in silico findings, we performed dose-escalation proliferation assays. Across different cell lines, ETS1-overexpressing cells exhibited increased sensitivity to the HSP90 inhibitor Alvespimycin (**Figure 4B**), but not to the HMGCR inhibitor Simvastatin or the Na⁺/K⁺-ATPase inhibitor Resibufogenin (**Supplementary Figure 3A**). We further confirmed these results using another HSP90 inhibitor, STA9090, in both TE5 and TT ETS1-overexpressing cells (**Figure 4B**). In parallel, colony formation assays performed with Alvespimycin and STA9090 consistently demonstrated enhanced sensitivity of ETS1-overexpressing cells to these HSP90 inhibitors (**Figure 4C**), whereas ETS1 knockdown in KYSE150 and HSC3 cells conferred reduced sensitivity (**Figure 4D; Supplementary Figure 3B**).

**Figure 4.**
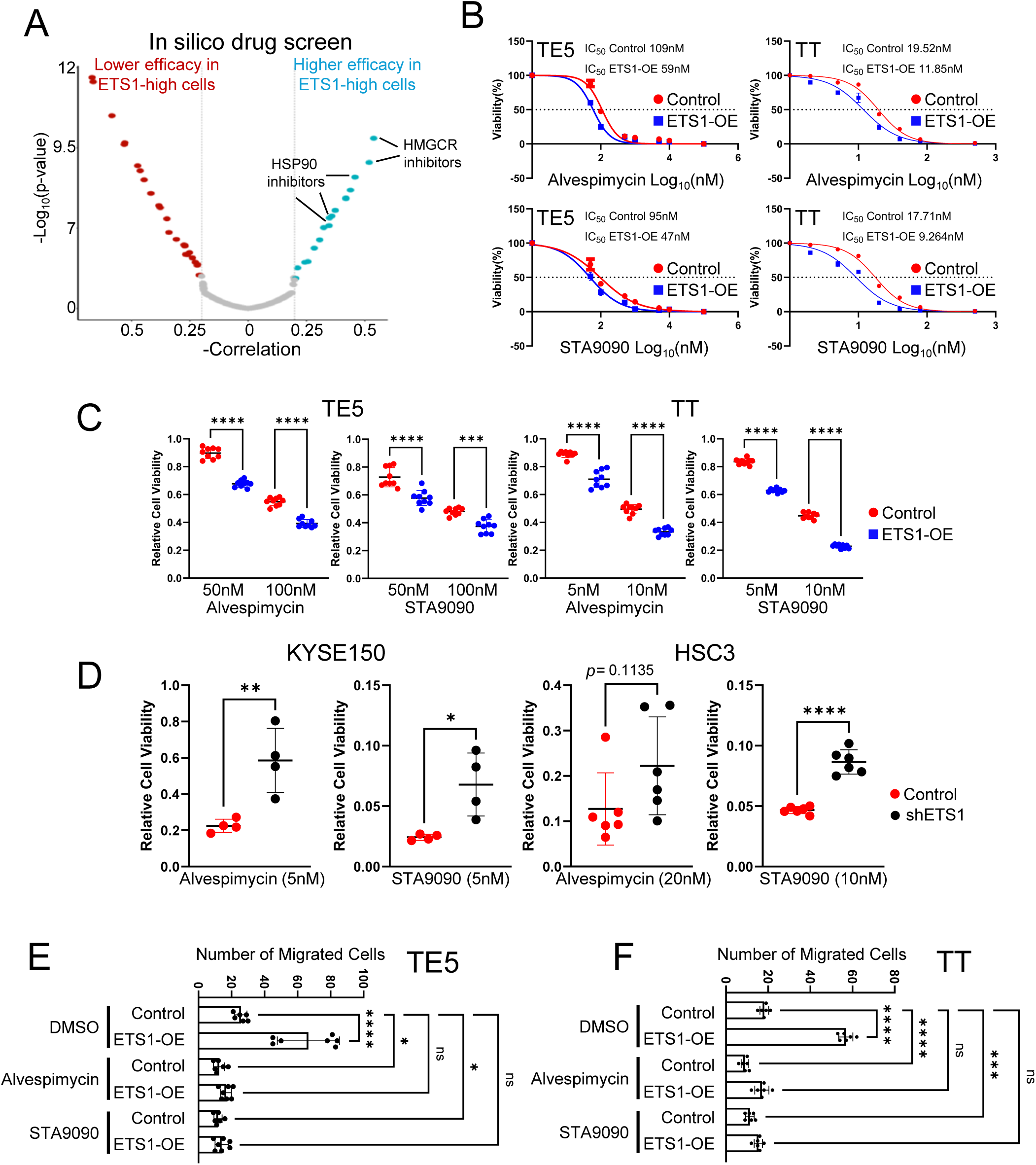
HSP90 inhibitors selectively target ETS1-high cancers. (A) Volcano plot showing in silico drug screen results based on the PRISM Primary Repurposing dataset^24^. (B) Cell viability (MTT) IC50 assays in both control and ETS1-overexpressing UASCC cell lines. Compounds are represented in nM using the Log10 scale. Data points represent the mean ± SD of at least two experimental replicates. (C-D) Colony formation assays upon exposure to Alvespimycin or STA9090 in control, ETS1-overexpressing, and ETS1-knockdown UASCC cell lines. Dot plots represent the mean ± SD of three experimental replicates. *P<0.05, **P<0.01, ***P<0.001, ****P<0.0001; P-values were determined using an unpaired t-test. (E-F) Transwell migration assays in control and ETS1-overexpressing UASCC cell lines with or without the exposure to Alvespimycin or STA9090. Bar graphs show the number of migrated cells and represent the mean ± SD of three experimental replicates. *P<0.05, ***P<0.001, ****P<0.0001; P-values were determined using a one-way ANOVA with multiple comparisons.

Based on our findings that HSP90 inhibitors modulate ETS1 expression, we next investigated whether these agents could reverse the migration properties driven by ETS1. Treatment with either Alvespimycin or STA9090 significantly reduced cell migration in both wound healing and transwell migration assays, effectively counteracting the enhanced migratory phenotype induced by ETS1 overexpression (**Figure 4E-F; Supplementary Figure 4A-B**). Conversely, knockdown of ETS1 resulted in the opposite results in these assays (**Supplementary Figure 4C**).

To elucidate how HSP90 inhibitors modulate ETS1, we treated cells with low doses of Alvespimycin and STA9090, and assessed ETS1 expression at both the mRNA and protein levels. Notably, both compounds effectively reduced the expression of ETS1 (**Figures 5A-B**). Seeking to understand the underlying mechanism, we considered prior studies identifying HIF1A as a direct client of HSP90^25^, and noting that HIF1A was reported to bind the ETS1 promoter in glioma cells^26^. Analysis of publicly available HIF1A ChIP-seq datasets across multiple tissues confirmed that HIF1A occupies the promoter region of ETS1 (**Figure 5C**). We validated these findings through ChIP-qPCR assays in control and ETS1-overexpressing cells, demonstrating HIF1A enrichment at the ETS1 promoter (**Figure 5D**). As a positive control for the ChIP-qPCR assay, we included ETS1 self-binding, a known feature of this transcription factor. To functionally confirm the regulatory relationship, we silenced HIF1A using siRNA, which led to significant reductions in ETS1 mRNA and protein levels across multiple cell lines (**Figures 5E-F**). Interestingly, HSP90 inhibitors also reduced levels of ectopically expressed ETS1 protein (**Figures 5G-H**), suggesting that post-translational mechanisms may contribute to ETS1 suppression. Together, these findings indicate that HSP90 inhibitors downregulate ETS1 at multiple levels, at least in part through disruption of HIF1A-mediated transcriptional activation.

**Figure 5.**
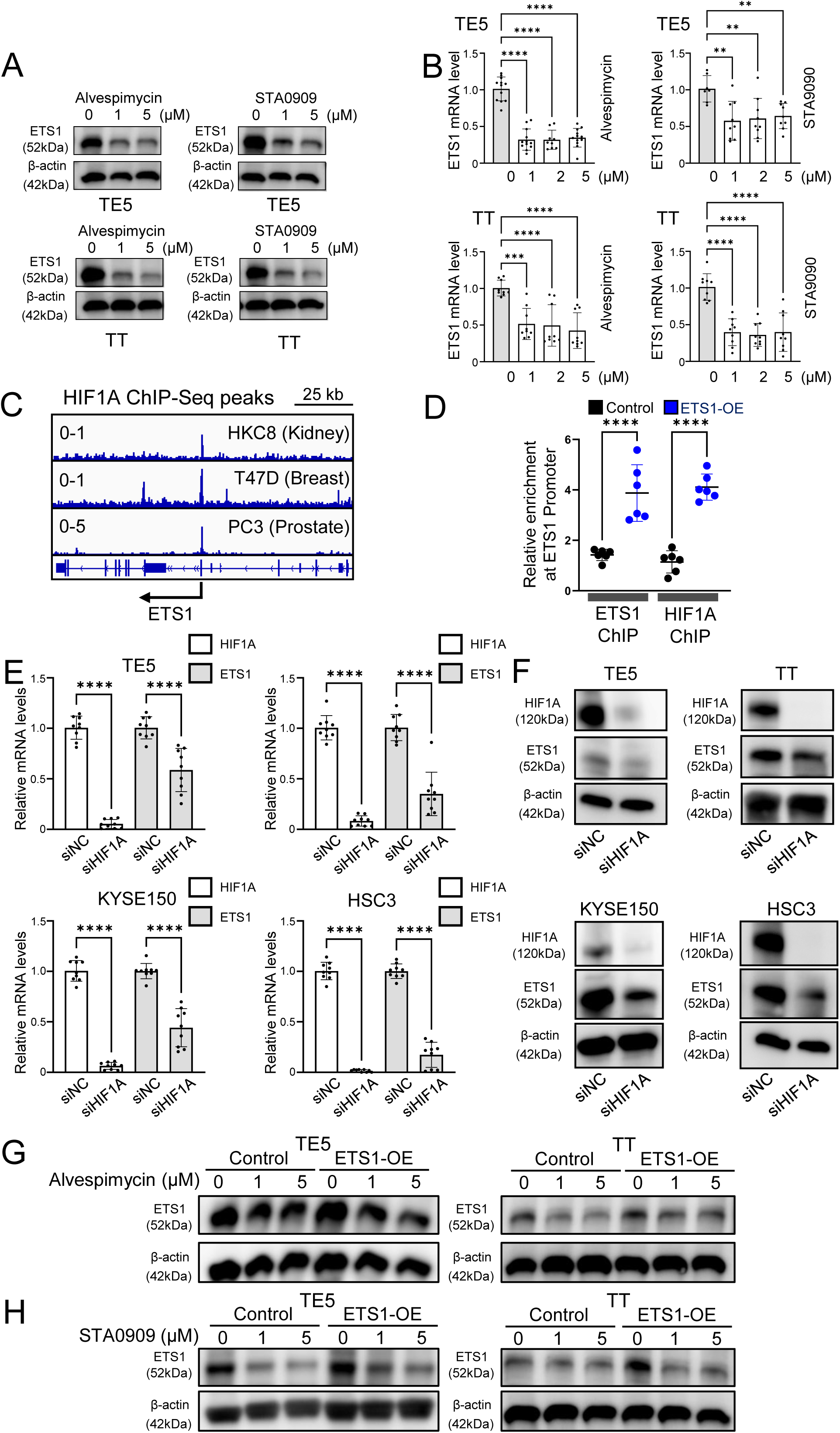
Identification of HIF1A as a key upstream regulator of ETS1. (A-B) The protein (A) and mRNA (B) levels of ETS1 in UASCC cell lines treated with Alvespimycin and STA9090 for 24 hours. Bar graphs represent the mean ± SD of three experimental replicates. **P<0.01, ***P<0.001, ****P<0.0001; P-values were determined using a one-way ANOVA with multiple comparisons. Western blot images are representative of three experimental replicates. (C) IGV plots of HIF1A ChIP-Seq in indicated samples at the ETS1 promoter region. Signal values of normalized peak intensity are shown in the upper left corner. (D) ChIP-qPCR assays measuring ETS1 and HIF1A binding on the promoter of ETS1 in TE5 control (black) and ETS1 overexpressing (blue) cells. Data is represented as fold change relative to IgG control. Individual data points from two replicates are shown. ****P<0.0001; P-values were determined using a one-way ANOVA with multiple comparisons. (E-F) HIF1A and ETS1 mRNA expression (E) and protein (F) levels in UASCC parental cell lines treated with siNC or siHIF1A for 48 hours. Bar graphs represent the mean ± SD of three experimental replicates. ****P<0.0001; P-values were determined using a one-way ANOVA with multiple comparisons. Western blot images are representative of three experimental replicates. (G-H) The protein levels of ETS1 in control and ETS1-overexpressing UASCC cell lines with or without the exposure to Alvespimycin or STA9090 for 24 hours. Western blot images are representative of three experimental replicates.

### ETS1 regulates the tumor microenvironment of UASCC

To further investigate the broader role of ETS1 as a transcriptional regulator, we performed RNA-Seq comparing control and ETS1-overexpressing (ETS1-OE) TE5 cells. Gene set enrichment analysis using the Hallmark gene sets revealed that the EMT pathway, as anticipated, was the most significantly upregulated program in ETS1-OE cells. Unexpectedly, several immune-associated signaling pathways, including TNFα and interferon pathways, were also significantly upregulated (**Figure 6A**). This is noteworthy, as ETS1 has not been previously implicated in the regulation of anti-tumor immune response. These findings are consistent with emerging evidence highlighting the interplay between metastatic processes and immune modulation within the tumor microenvironment (TME)^27,28^. Corroborating our experimental data, analysis of ETS1-high tumors in TCGA HNSCC and ESCC cohorts similarly demonstrated significant enrichment of immune-related gene signatures (**Figure 6B**).

**Figure 6.**
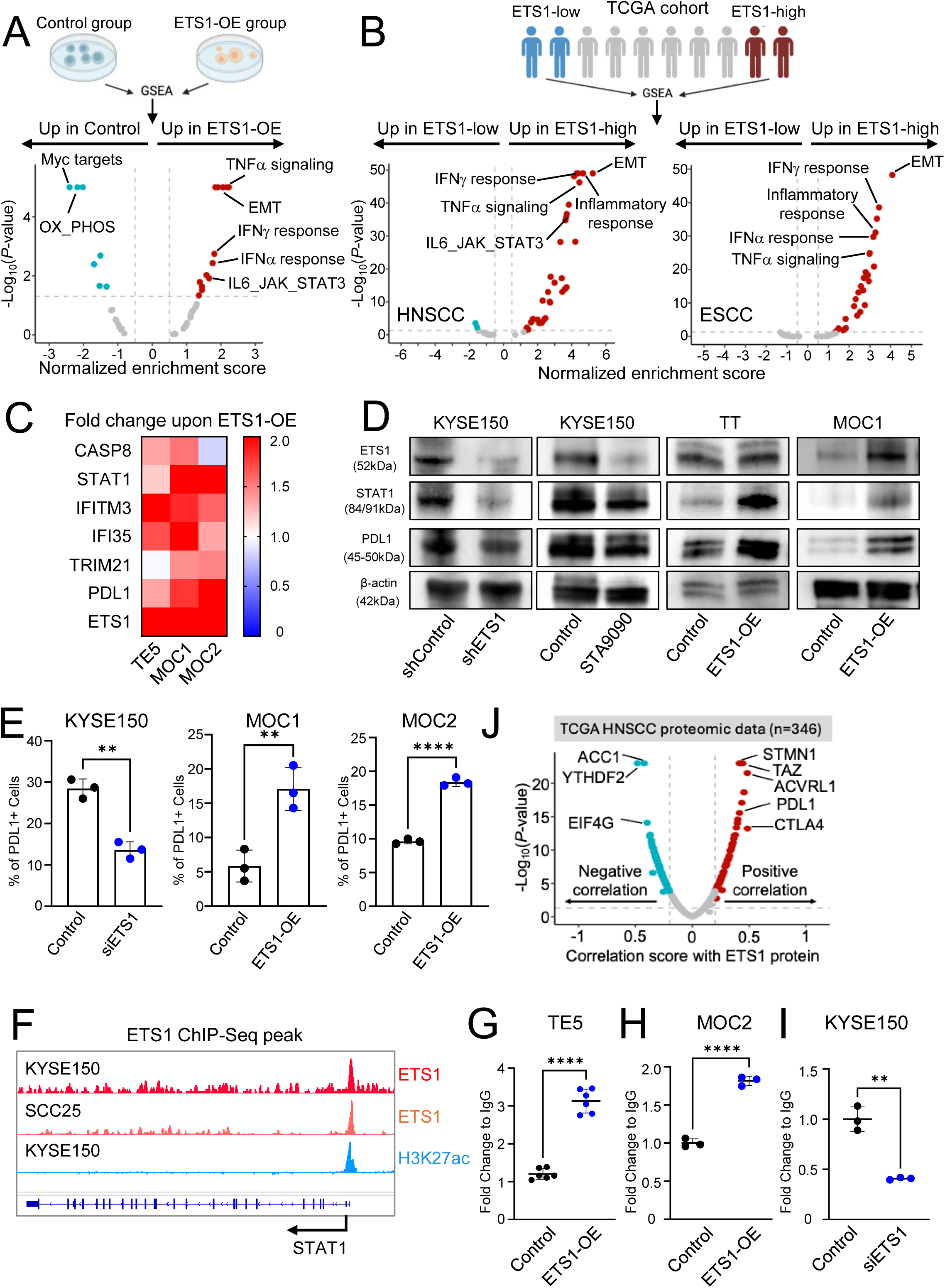
ETS1 regulates transcription of STAT1 and PDL1. (A–B) Volcano plots showing GSEA results of enriched Hallmark pathways in (A) ETS1-overexpressing vs. control TE5 cells and (B) ETS1-high vs. ETS1-low tumor samples from TCGA HNSCC and ESCC cohorts. Significantly enriched pathways are highlighted in color. (C) Heatmap showing the fold change in expression of selected leading-edge genes in ETS1-overexpressing vs. control UASCC cells. Data represent the mean of three experimental replicates. Expression levels range from high (red) to low (blue), with values at or above a fold change of 2 displayed as the maximum (red). (D) Representative western blot images showing protein levels of STAT1, PDL1, and ETS1 in UASCC cells under the indicated conditions. Images are representative of three independent experiments. (E) Quantification of flow cytometry analysis of PDL1 cell surface expression. **P<0.01, ****P<0.0001; P-values were determined using an unpaired t-test. (F) IGV plot depicting ETS1 ChIP-Seq signal at the STAT1 promoter region in KYSE150 cells. Normalized peak intensity values are indicated in the upper left corner. (G–I) ChIP-qPCR analysis of ETS1 binding at the STAT1 promoter in the indicated cell lines. Data are presented as fold enrichment relative to IgG control. Individual data points from two replicates are shown. ***P<0.001, ****P<0.0001; P-values were determined using an unpaired t-test. (J) Volcano plot showing proteins correlated with ETS1 in TCGA HNSCC proteomics data. Significantly correlated proteins are highlighted in color.

To validate these findings, we examined a subset of genes from the immune-related pathways enriched in ETS1-OE cells. Consistently, we observed significant upregulation of these genes across multiple cell lines (**Figure 6C**), supporting a role for ETS1 in modulating immune response signaling. Among these genes were STAT1, a central regulator of immune responses, and PD-L1 (CD274), a key immune checkpoint molecule with a fundamental role in anti-tumor immunity. Notably, PD-L1 is an FDA-approved therapeutic target in human cancers. Given their biological and clinical relevance, we further validated ETS1-mediated regulation of STAT1 and PD-L1 expression by Western blotting in multiple cell lines (**Figure 6D**). We also confirmed PD-L1 upregulation at the cell surface using flow cytometry analysis (**Figure 6E**).

With respect to STAT1, we found that ETS1 directly occupies the promoter region of the STAT1 gene in both human and murine samples, as demonstrated by ChIP-Seq (**Figure 6F**) and further validated by ChIP-qPCR (**Figures 6G-I**). In addition, analysis of TCGA HNSCC proteomics data revealed that PD-L1 protein abundance is among the most significantly positively correlated with ETS1 protein levels (**Figure 6J**). Collectively, these results highlight a regulatory link between ETS1 and STAT1/PD-L1, implicating ETS1 as a potential modulator of key anti-tumor immune regulators.

Given the observed regulation of immune modulators by ETS1, we next investigated the functional consequences of ETS1 overexpression on anti-tumor immune responses in vivo. To this end, we employed the syngeneic MOC2 model and established allografts in immunocompetent C57BL/6J mice. To assess immune cell composition within the TME, we performed flow cytometry analysis on dissociated tumors. Notably, ETS1-overexpressing tumors exhibited significantly reduced infiltration of CD3⁺ T cells (**Figure 7A**), including both CD4⁺ (**Figure 7B**) and CD8⁺ (**Figure 7C**) T cell subsets, compared with control tumors. Consistent with these findings, we also observed a decreased abundance of T cells in cervical lymph node samples from the ETS1-overexpressing group (**Figure 7D**). We further validated these findings using immunohistochemistry (IHC) staining, which confirmed a marked reduction in CD3^+^ and CD8⁺ T cell infiltration within ETS1-overexpressing tumors compared to control tumors (**Figure 7G-H**). Importantly, ETS1 overexpression significantly increased the rate of tumor growth (**Figure 7E**) and markedly reduced overall survival in tumor-bearing mice (**Figure 7F**), suggesting that ETS1 may promote immune evasion and tumor progression through suppression of anti-tumor T cell responses.

**Figure 7.**
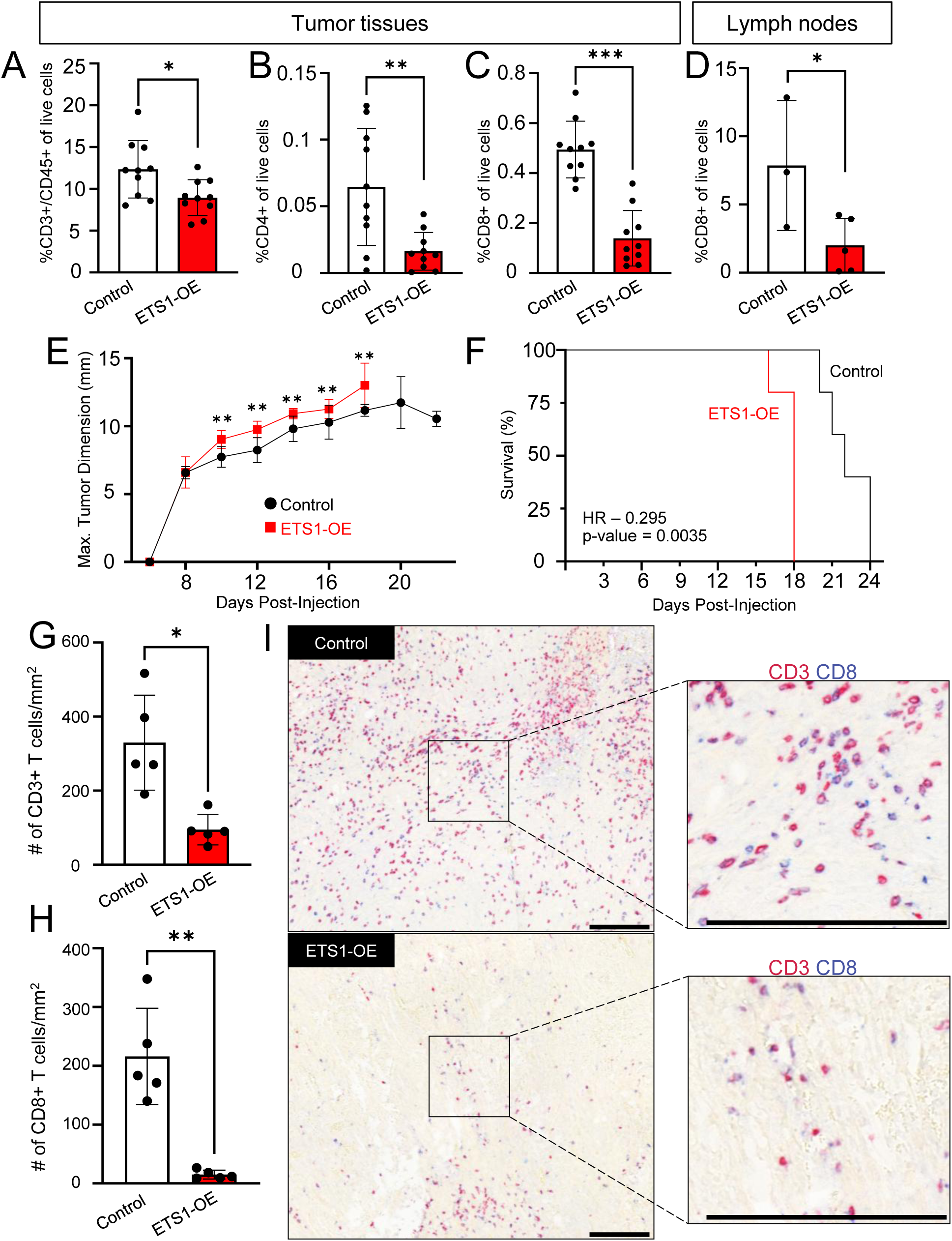
ETS1 overexpression impairs T cell immunity and promotes tumor growth. (A–D) The proportion of CD3^+^, CD4^+^ and CD8^+^ T cells out of total live cells from control or ETS1-overexpressing (A-C) allografts, or (D) cervical lymph nodes, measured by flow cytometry analysis. Bars represent mean ± SD of independent samples. *P<0.05, **P<0.01, ***P<0.001; P-values were determined using an unpaired t-test. (E-F) Allografts growth curves and survival curves of control or ETS1-overexpressing tumor-bearing mice. *P<0.05, P-values were determined using a two-way ANOVA (E). P-value was determined by the log-rank test (F). (G-I) IHC of control and ETS1-overexpressing tumors stained with CD3^+^ T cells (red) and CD8^+^ T cells (blue). Quantification of CD3^+^ T cells (G) and CD8^+^ T cells (H) are provided. (I) Images are representative of 10 tumors, 5 control and 5 ETS1-overexpressing. Scale bar = 0.1mm.

To determine whether ETS1 overexpression impairs CD8⁺ T cell-mediated tumor killing, we established an *ex vivo* co-culture assay. MOC2 cells were pulsed with the OVA peptide (SIINFEKL), allowing surface presentation via MHC-I, then washed and co-cultured with OT-I CD8⁺ T cells, which specifically recognize OVA-MHC-I complexes (**Figure 8A**). At a 10:1 effector-to-target ratio, OT-I T cells eliminated ∼40% of control tumor cells (**Figure 8B**). Importantly, ETS1 overexpression significantly reduced this antigen-specific cytotoxicity (**Figure 8B**), indicating that ETS1 impairs tumoricidal activity of CD8⁺ T cells in UASCC cells.

**Figure 8.**
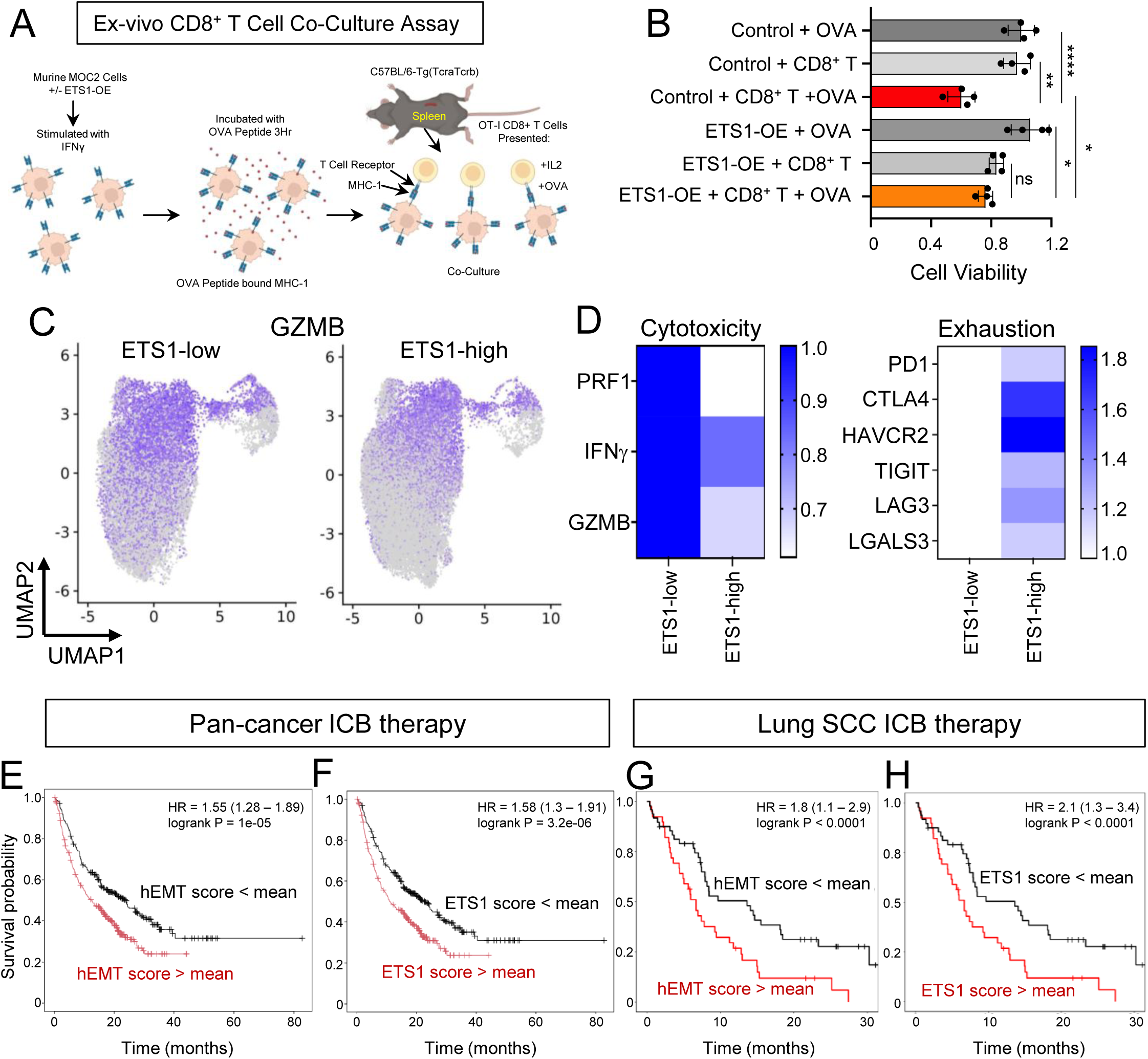
ETS1 expression is associated with poor immunotherapy response. (A) A schematic graph showing experimental design of ex vivo CD8^+^ T cell co-culture assay. (B) Bar graphs showing relative cell viability of MOC2 cells co-cultured with CD8^+^ T cells as described in (A). Bars show the means ± SD representative of two 4 replicates. *P < 0.01, **P < 0.01, ***P < 0.001. P-values were determined using unpaired t-tests. (C-D) UMAP plots showing the expression levels of GZMB and (D) quantification of T cell cytotoxic or exhaustive marker genes of CD8^+^ T cells from ETS1-high vs. -low HNSCC tumors. The average ETS1 expression in cancer cells in each tumor was calculated and used for sample classification. (E-F) Patient survival curves analyzed using the pan-cancer ICB-treated cohort (n=976 patients across multiple cancer types^29^) via the KM plotter^41,42^. Patients were stratified using either the mean hEMT score (E) or mean ETS1 score (F). (G-H) Patient survival curves analyzed using a lung squamous carcinoma cohort treated with anti-PDL1 antibody (n=87)^30^. Patients were stratified using either the mean hEMT score (G) or mean ETS1 score (H).

### ETS1-signature is associated with ICB resistance in human ICB clinical outcome

Our data suggest that ETS1 drives an immune-cold TME in UASCC by activating both STAT1 and PDL1. To explore the translational relevance of this regulatory pathway, we analyzed the earlier combined scRNA-seq data from 114 UASCC patient samples^4,17–20^. Specifically, we stratified tumor samples into ETS1-high and ETS1-low groups (top and bottom 20%, respectively) based on ETS1 expression levels in cancer cells. Notably, the inferred activity of CD8^+^ T cells, as characterized by the expression of cytotoxic markers such as GZMB, PRF1, and GZMK, was elevated in ETS1-low tumors (**Figures 8C-D**). Conversely, ETS1-high tumors had upregulation of multiple T cell exhaustion markers, including PD1, CTLA4, HAVCR2, TIGIT, LAG3, and LGALS3 (**Figure 8D**). These findings highlight a potential link between ETS1 expression, immune evasion mechanisms, and the suppression of anti-tumor T cell responses in UASCC.

Encouraged by these findings, we next investigated whether ETS1 contributes to immune checkpoint blockade (ICB) therapy response. Given that ETS1 is also expressed in subsets of lymphocytes, we derived an ETS1-gene signature to infer ETS1 activity. This signature was developed using two approaches: the 50 hEMT gene set (termed the “hEMT score”) and the 35 ETS1-binding target genes (termed the “ETS1 score”) identified in our earlier analyses (**Figures 1A, 2F**). To assess the clinical relevance of these gene signatures, we first analyzed a pan-cancer ICB-treated cohort comprising 976 patients across multiple cancer types^29^. Notably, both ETS1-gene signatures were strongly associated with resistance to ICB therapy, as evidenced by poorer clinical outcomes in signature-high patients (**Figures 8E-F**). To further validate these observations in a squamous cancer-specific context, we examined an independent lung squamous carcinoma cohort in which patients were treated with anti-PD-L1 therapy^30^. Notably, low ETS1-signatures were significantly associated with improved survival in ICB treatment (**Figures 8G-H**). These concordant findings across multiple independent clinical cohorts highlight the predictive value of ETS1 activity for determining ICB therapy response. Collectively, our results suggest that ETS1 expression and activity may serve as a biomarker for resistance to immunotherapy, providing a potential tool for patient stratification and treatment optimization in the clinical management of UASCC and related malignancies.

## Discussion

Each tumor contains diverse transcriptional and cellular states that underlie ITH, a major driver of tumor metastasis and disease recurrence—making it a critical barrier to effective clinical management^3^. At the scientific level, transcriptional ITH represents a fundamental challenge in understanding cancer biology, tumor evolution, and therapeutic resistance. Our scRNA-seq analysis revealed an hEMT program as a critical mediator of transcriptional ITH and metastatic dissemination in UASCC. While canonical EMT is well-studied in cancer progression, the hEMT state—marked by co-expression of epithelial (e.g., CDH1, IRF6, JUP) and mesenchymal (LAMC2, ITGB4, TGFA) genes—has emerged as a driver that enhances cellular plasticity and tumor metastasis. Our findings align with recent work in HNSCC showing that the hEMT state is strongly associated with invasion and migration^4,5,9^.

Although ETS1 has been implicated in classical EMT, its role in maintaining hEMT has not been investigated. ETS1’s ability to simultaneously activate mesenchymal genes and suppress full epithelial dissolution suggests it stabilizes a metastable hEMT state. This duality may explain why ETS1-high tumors exhibit aggressive dissemination in vivo, a finding not recapitulated in prior in vitro cell line studies (e.g., CAL27/SCC25 knockdown models^21,31^). Our in vivo metastasis assays and clinical cohort analyses suggest ETS1’s role as a driver of progression, bridging the gap between cell-autonomous EMT and systemic spread.

A translational insight from our study is the sensitivity of ETS1-high tumors to HSP90 inhibitors. HSP90 is a chaperon protein with hundreds of protein substrates involved in the folding of proteins. Among these client proteins, it has been found that HSP90 can regulate the hypoxia inducible factor 1 alpha (HIF1A)^25,32,33^. Mechanistically, we traced this vulnerability to HIF1A suppression: HSP90 stabilizes HIF1A, which binds the ETS1 promoter to sustain its expression. Such a pathway is in agreement with prior findings^26,34^ wherein HIF1A (possibly induced by tumor hypoxia) amplifies ETS1, which in turn promotes metastatic traits. Translationally, this dependency is actionable—HSP90 inhibitors like Alvespimycin and STA9090 (ganetespib) reduced ETS1 levels and reversed its pro-metastatic effects. Although early-phase trials^35^ of HSP90 inhibitors showed limited efficacy, our work suggests biomarker-driven patient selection (e.g., ETS1-high tumors) could revive this therapeutic strategy.

Beyond metastasis, we uncovered a previously underappreciated role for ETS1 in anti-tumor immune regulation. ETS1-high tumors exhibited suppressed T cell infiltration and upregulated PD-L1, a phenotype mediated by ETS1’s direct activation of STAT1. STAT1, often associated with pro-inflammatory signaling, paradoxically drove immune suppression, likely by amplifying PD-L1 and other checkpoint molecules (CTLA4, LAG3). This aligns with evidence that STAT1 can polarize tumors toward immune exclusion^36^. Notably, ETS1’s immune-regulatory function appears to be distinct from its roles in NK or T cell development^37,38^, highlighting context-dependent effects. The link between ETS1 and immune evasion has direct clinical relevance. In multiple ICB-treated cohorts, ETS1-high tumors were resistant to the ICB therapy, likely due to their immune-cold phenotype. Our ETS1 gene signature (derived from hEMT and ChIP-seq targets) could stratify patients likely to fail the ICB therapy, guiding alternative strategies such as combination approaches integrating HSP90 inhibition, or agents that remodel the TME to restore immune infiltration and thereby sensitize tumors to immunotherapy. In support, our recent work in HNSCC demonstrates that HSP90 inhibitors enhance T cell anti-tumor immunity in murine syngeneic models^39^.

In summary, our study establishes ETS1 as a central driver in UASCC progression, integrating key biological processes that promote metastasis through hEMT, drive immune evasion via activation of expression of both STAT1 and PD-L1, and confer resistance to immune checkpoint blockade. These findings not only provide mechanistic insights into the multifaceted role of ETS1 in tumor biology but also highlight its potential as a biomarker and therapeutic target. Going forward, several important questions remain to be addressed. For example, does the function of ETS1 in promoting hEMT depend on specific tumor spatial contexts, such as hypoxic regions? Can therapeutic strategies that combine HSP90 inhibition with immune checkpoint blockade synergistically overcome ETS1-mediated immune resistance and improve clinical outcomes? Future studies integrating spatial transcriptomics, functional organoid models, and preclinical therapeutic testing will be essential to address these questions and translate our findings into effective therapeutic interventions for UASCC and related malignancies.

## Acknowledgments

This work was supported by the NIH under the award R37CA237022, R01DE033648, 1R01DK135562 as well as P30CA014089 (D.-C.L.), and Watt Family Endowed Chair for Head and Neck Cancer Research (U.K.S.).

## Materials and Methods

### Human and mouse cell lines

Human UASCC cancer cell lines HSC3, TE5, and KYSE150 cells were grown in RPMI-1640 medium (Corning, 45000-396). TT cells were cultured in Dulbecco’s modified Eagle medium/Hams F-12 50:50 (DMEM:F12) (Corning, 45000-344). Both media were supplemented with 10% FBS (Omega Scientific, FB-02) and 1% penicillin-streptomycin sulfate (Gibco, 10378016). Murine oral SCC cells, MOC1 and MOC2, were cultured in Iscove’s Modified Dulbeccos Medium (IMDM) (HyClone, SH30228.01), with 10% heat-inactivated fetal bovine serum (Omega Scientific, FB-02) and 1% penicillin-streptomycin sulfate (Gibco, 10378016). All cultures were maintained at 37°C supplemented and with 5% CO2. Cell lines were regularly tested for mycoplasma and verified by us using short tandem repeat analysis.

### Cell proliferation and colony formation assays

Cells were seeded in 50uL of media into 96-well plates (1,000–5,000 cells/well) in quadruplicates. Alvespimycin (Selleckchem, S1142), STA9090 (Selleckchem, S1159), Simvastatin (Selleckchem, S1792), Resbufogenin (Selleckchem, S3238), or DMSO (VWR, 97063-136) was then mixed into the wells containing the seeded cells. After 72 hours, cell proliferation was measured using 10uL of MTT (3-(4, 5-dimethylthiazol-2-yl)-2, 5-diphenyl tetrazolium bromide) (Sigma, #475989). Crystals were dissolved by incubating plates overnight with the 100uL of 10% SDS, 0.01M HCl. Plate absorbance readings were taken using a 570nm wavelength. For colony formation assays, cells were seeded into six-well plates (1,000–3,000 cells/well) overnight. On the following day, the media was replaced with fresh media containing either Alvespimycin (Selleckchem, S1142), STA9090 (Selleckchem, S1159), or DMSO (VWR, 97063-136), and cultured for ∼2 weeks. The colonies were then fixed using 4% paraformaldehyde (Electron Microscopy Sciences, 15711) and stained with 1% crystal violet (Sigma, V5265). Colonies were further dissolved using 100uL of 10% SDS and the absorbance readings were detected at a 595nm wavelength.

### RNA extraction, cDNA synthesis, and quantitative PCR

Total RNA was extracted using the RNeasy Mini kit (QIAGEN, 70106) and cDNA was reverse transcribed from the total RNA using the LunaScript RT SuperMix cDNA Synthesis kit (New England BioLabs, M3030L). Quantitative PCR (qRT-PCR) was conducted with PowerUp™ SYBR™ Green Master Mix (Thermo Scientific, A25918). Primers used in this study are listed in Supplementary Table 2.

### Chromatin immunoprecipitation (ChIP)

ChIP was performed as described previously^43^. Approximately 1 × 10L cells were fixed in 10mL of fresh media containing 1% paraformaldehyde (Electron Microscopy Science, 15710-S) for 10 minutes at room temperature, then quenched with 1ml of 1.25M glycine for 5 minutes and washed 3 times with ice cold 1 x PBS. Cells were collected by centrifugation at 1,500 x g for 5 minutes. The supernatant was removed, and cells were lysed twice with 1 ml lysis buffer (formula: 150mM NaCl, 5 mM EDTA pH 7.5, 50mM Tris pH 7.5, 0.5% NP-40) containing protease inhibitors (Roche, 04693124001). Cells were first lysed mechanically by pipetting in a microcentrifuge tube and incubated on ice for 5 minutes. Next, cells were passed through a 23G/1mL syringe, five times, and incubated on ice for 5 minutes. Cells were then spun down at 9,400 × g for 5 minutes at 4 °C and the supernatant was removed. This lysis process was repeated using 1mL of lysis buffer. Cell pellets were then resuspended in 1mL shearing buffer (formula: 1% SDS, 10mM EDTA pH 8.0, 50 mM Tris pH 8.0) and transferred into 15mL conical tubes. Cells were sonicated using a Diagenode Bioruptor sonicator with 130 mg of sonication beads for 15 cycles (30 seconds on, 30 seconds off). Next, sonicated cells were transferred into microcentrifuge tubes and spun down at 20,000 × g for 10 min at 4 °C. The supernatants were collected into a new microcentrifuge tube and a 10uL aliquot of the supernatants was set aside and stored at −20C to be used as the input sample, while the remaining supernatant was further diluted 1:5 in dilution buffer (formula: 1.1% Triton X-100, 0.1% SDS, 1.2mM EDTA pH 8.0, 167 mM NaCl, 16.7mM Tris pH 8.0). 2 μg ETS1 antibody (Cell Signaling, 14069) or HIF1A antibody (Cell Signaling, 36169) was then added and incubated at 4 °C overnight on a rotator. On the next day, 30uL of Dynabeads Protein G beads (Thermo Scientific, 10004D) were carefully added to the samples and incubated for 4 hours on a rotator at 4 °C. Dynabeads were then separated using a magnet and washed 10 times with ice cold lysis buffer, followed by two washes with ice cold TE buffer (formula: 10mM Tris pH 8.0, 1 mM EDTA pH 7.5, adjusted to a final pH of 7.6). DNA samples were released from beads using 100uL of reverse crosslinking buffer (formula: 136mM NaHCO3, 0.96% SDS) while rotating at room temperature for 15 minutes, repeated twice. The DNA samples were pooled together into a PCR tube containing 4uL of 5M NaCl and incubated for 14-16 hours at 65 °C. Next, 1uL of RNase (Thermos Scientific, # EN0531) was mixed into samples and further incubated at 37 °C for 30 minutes. Then 4uL 0.5M EDTA pH 8.0 with 8uL of 1M Tris pH 7.0, and 1uL of Proteinase K (10mg/mL) (Invitrogen, #100005393) was mixed into the sample and incubated for 1 hour at 45°C. Purification of DNA was performed by adding 500uL of Buffer PB (Qiagen, 19066) to samples, vortexing, and transferring samples into a spin column. Samples were incubated for 2 minutes at room temperature. Samples were spun down at 17,900g for 2 minutes, and the supernatant was discarded. The columns were then washed with Buffer PE (Qiagen, 19065) three times. The DNA was eluted by adding 50uL of sterile water to the columns and incubated for 5 minutes before spinning down columns at 17,900 x g for 2 minutes. The purified DNA was used for either quantitative PCR or DNA library preparation and deep sequencing using Illumina HiSeq platform.

### Western blotting

Cells were lysed using the RIPA lysis buffer system (Santa Cruz Biotechnologies, sc-24948A), supplemented with proteinase inhibitor cocktail, PMSF, and phosphatase inhibitors for 30 min on ice. Protein concentrations were quantified using the Bradford reagent (VWR, E530-1L). Western blotting was performed as previously described using SDS-PAGE gels (GenScript, #M41210) and transferred to 0.45um immune-blot PVDF membrane (Millipore, #IPVH00010) for 60 minutes at 100V using the transfer buffer (formula: 25mM Tris, 192mM Glycine, 20% Methanol)^44^. Membranes were then blocked, and primary antibodies were diluted in TBST (formula: 20mM Tris, pH 7.4, 140mM NaCl, 0.1% Tween-20) with 5% BSA (Research Products International, #9048-46-8). Primary antibodies were incubated overnight at 4°C for 14-16 hours. Membranes were washed in TBST for 5 minutes, a total of 4 times, and then incubated with secondary antibodies for 1 hour at room temperature in TBST-SDS (formula: 20mM Tris, pH 7.4, 140mM NaCl, 0.2% Tween-20, 0.01% SDS) with 5% non-fat powdered milk (Bioworld, #30620074-1). Membranes were washed in TBST for 5 minutes, a total of 4 times, and then twice with 1X PBS. Membranes were developed using an ECL substrate (Thermoscientific, A38554) and chemiluminescence exposures. Primary antibodies used were ETS1 (1:1,000, Cell Signaling, 14069), HIF1A (1:1,000) (Cell Signaling, 36169), STAT1 (1:1,000, Cell Signaling, 14994), PD-L1 (1:1000, Proteintech, #66248-1-Ig) and Actin (1:3,000, Developmental Studies Hybridoma Bank, JLA20). Secondary antibodies were goat anti-rabbit HRP (Millipore Sigma, 112-348) and goat anti-mouse HRP (Invitrogen, G21040).

### Transfections and lentiviral production

Cells were transfected with non-silencing siRNA or siHIF1A (IDT, hs.Ri.HIF1A.13) using Lipofectomine RNAiMAX (Thermo Fisher, #13778150). Lentiviral cloning vector pLKO.1-TRC (Addgene, #10878) was used for shRNA expression. The double-stranded oligonucleotide shRNAs for human and mouse ETS1 sequences were ligated into the AgeI/EcoRI sites of the pLKO.1-TRC digested lentiviral vector. shRNA target sequences are listed in Supplementary Table 2. Transfection of second-generation lentiviral vectors included 2ug of lentiviral vector (pLKO.1 TRC), packaging vectors pMD2.G 0.5ug (Addgene, #12259) and pPAX2 1.5ug (Addgene, #12260), using 100uL of serum free DMEM media (Corning, 45000-304) and 6uL of BioT lipofectamine (Bioland Scientific, #B01-00). For overexpression cloning, ETS1 overexpression CMV vectors were purchased from the Vectorbuilder. pLenti CMV V5-LUC Blast vector was from Addgene (#21474). Transfection of third generation lentiviral vectors included 2ug of lentiviral vectors packaging vectors pRSV-Rev 1ug (Addgene, #12253), pMDLg/pRRE 1.5ug (Addgene, #12251), and pMD2.G 0.5ug (Addgene, #12259) using 100uL of serum free DMEM media (Corning, 45000-304) and 6uL of BioT lipofectamine. The recombinant viral vectors and packaging vectors were then co-transfected into HEK293T cells. Supernatants were passed through a 0.45 μM syringe filter 48 hours after transfection. Target cells were infected with the virus in the presence of 5 ug/ml Polybrene (Santa Cruz Biotechnology, #134220).

### Selection of stable knockdown and overexpression cell lines

Upon infection with viral particles, cells were selected using puromycin (Santa Cruz Biotechnology, #205821). Luciferase infected cells were selected using Blasticidin. Stable knockdown and overexpression cells remained in media containing puromycin throughout the length of culture, and knockdown/overexpression was confirmed using quantitative PCR/WB prior to experiments.

### Xenograft and *In Vivo* studies

Animal studies were performed in accordance with protocols approved by the ethical regulations of the Institutional Animal Care and Use Committee (IACUC) of USC. To generate syngeneic orthotopic tumors, 2.5 × 10^5^ MOC2 cells were mixed 1:1 with BME (Cultrex, #3533-005-002) to a final volume of 100uL and subcutaneously injected into the right cheek of six-week-old C57BL/6 mice. Tumors were serially measured with electronic calipers. Tumor volume was calculated with the formula V = (Width X Height^2^)/2. Tumors were allowed to grow until they reached a maximum of 12mm in any direction or mice experienced significant tumor-associated morbidity, at which point mice were euthanized and tumors and lymph nodes were collected for further studies.

For tail vein injection studies, 1 × 10^6^ KYSE150 (scramble or shETS1) or 2 × 10^6^ TE5 (scramble or ETS1-overexpressing) cells were injected into the tail vein of NOD-SCID Gamma (NSG) mice (six weeks old). For TE5 cells, bioluminescence imaging was performed periodically over the course of the experiments. The mice’s general behavior were monitored, and mice were euthanized when appearances indicated physical abnormalities. At the end of the experiments, mice were sacrificed, and the tumor tissues were collected.

### Flow Cytometry

Tumors from the cheek and lymph nodes of C57BL/6 mice were digested in collagenase/Hyaluronidase (Stemcell Technologies, #07912) and Dispase II (Thermo Scientific, #17105041) in DMEM media (Corning, 45000-304) to obtain single cell suspensions. Single cells (1 x 10^6^) in 100uL were next incubated on a rocker with a FC blocker (BD, #553141) for 10 minutes. Cells were then incubated with the following antibodies: 1 test volume of Zombie Violet Fixable Dye (BioLegend, #423113), 0.1μg of CD3 conjugated PerCP/Cyanine5.5 (BioLegend, #100217), 0.1μg of CD4 conjugated FITC (BioLegend, #100405), 0.1μg of CD8 conjugated PE (BioLegend, #100707), 0.1μg of CD45 conjugated Brilliant Violet 605 (BioLegend, #103155) for 30 minutes in the dark on a rotator. Cells were centrifuged at 3,900 x g for 1 minute and finally resuspended in 1mL of cold 1X PBS. Flow cytometry analysis was conducted using the AttuneNXT machine. Populations of CD45^+^CD3^+^CD4^+^CD8^−^, CD45^+^CD3^+^CD4^−^CD8^+^ T cells were quantified and analyzed. Data was processed and analyzed using FlowJo software (BD bioscience, Ver10.10).

To measure cell surface levels of PDL1, cells were trypsinized to single cell suspension and aliquoted to 1 x 10^6^ cells in 100uL cPBS (2% FBS + 98%PBS). Single cells were next incubated on a rocker with a FC blocker (BD, #553141) for 10 minutes. Cells were then incubated with the following antibodies: 0.2ug of human PDL1 conjugated with APC (Cytek, 20-5984-T025), or murine PDL1 conjugated with PE-Cy7 (Cytek, 60-1243-U025) for 45 minutes in the dark on a rotator. Flow Cytometry analysis was performed using the same method.

### Trans-well migration and wound healing assays

In a 24 well plate, 400uL of culture media containing 10% FBS was added to the well. A 6.5mm insert with 8.0 μm polycarbonate membranes (COSTAR, #3422) was placed on top of the wells. A total of 20,000–50,000 cells were resuspended in 100uL of serum free media and placed into the insert. Cells were incubated for 48-72 hours and then counted using a hemocytometer. For drug exposure experiments, cells were placed into serum free media containing drugs right before being placed into the inserts. Additionally, the same concentration of drug was added to the bottom portion of the well in media containing 10% FBS. For wound healing assays, cells were seeded in 6 well plates. Once the cells had reached ∼70% confluence, a wound was created along the surface of the cells and allowed 24 hours to recover. Images were taken immediately after the wound was created and 24 hours following the wound formation. For drug exposure experiments for wound healing assays, drugs were added at the time of wound induction and remained for the 24-hour incubation period.

### Immunohistochemistry

Formalin-fixed and paraffin-embedded tissues were cut at 4 µm thickness using a microtome then deparaffinized with xylene and ethanol. Peroxidase blocking was performed using Peroxidazed 1 (BioCareMedical, PX968) for 5 minutes at room temperature. Antigen retrieval was performed at 121C for 5 minutes using Reveal Decloaker solution (BioCareMedical, RV1000) in an ARC Decloaking Chamber (BioCareMedical, DCARC0001). Staining of slides was performed on an IntelliPATH PLUS automated staining platform (BioCareMedical, IPPLS0001US). Slides were blocked with Rodent Block M (BioCareMedical, RBM961) for 30 minutes at room temperature. Slides were incubated with an anti-CD3 (1:100, BioRad, MCA1477) for 2 hours at room temperature and detected using the Rat-on-Mouse AP-Polymer system (BioCareMedical, RT518). Slides were then stained using the IntelliPATH Fast Red Chromogen Kit (BioCareMedical, IPK5017). Slides were then incubated with Denaturing Solution (BioCareMedical, DNS001) for 10 minutes. Slides were again blocked with Rodent Block M for 30 minutes at room temperature, followed by a 2-hour incubation at room temperature with anti-CD8a (1:100, Invitrogen, 14-0808-80). Detection was performed using the Rat-on-Mouse AP-Polymer system and then stained using the IntelliPATH Ferangi Blue Chromogen Kit (BioCareMedical, IPK5027). Washes were performed after each application using TBS Buffer (BioCareMedical, TWB945). Slides were imaged using the Pannoramic 250 Flash III Digital Scanner (3Dhistech,P250). CD3^+^ and CD8^+^ T cells were quantified using ImageJ and normalized to the area of the tumors.

### Ex vivo co-culture of CD8^+^ T cells and cancer cells

MOC2 control and MOC2 mETS1-OE cells were incubated in two 10cm dishes in media supplemented with 20 ng/mL IFNγ (Abcam, #9922) for 24Lhr. Media was then aspirated, cells were washed with PBS, and media was replaced. MOC2 cells were then pulsed with 1 ng/mL OVA peptide (Selleckchem, E2915) for 3Lhr. Cells were trypsinized (Gibco, #15400-054), counted, and plated at 1,500 cells/well in 200uL of Iscove’s Modified Dulbeccos Medium (IMDM) (HyClone, SH30228.01), with 10% heat-inactivated fetal bovine serum (Omega Scientific, FB-02), 1% penicillin-streptomycin sulfate (Gibco, 10378016), and 5ug/mL puromycin (Thermo Scientific A1113803). MOC2 cells were incubated overnight. Simultaneously, spleens were removed from C57BL/6-Tg (TcraTcrb) mice and splenocytes were treated with 1X Red Blood Cell lysis buffer (eBioscience, #00-4333-57) for 10 min. Splenocytes were washed in 1X PBS and plated in RPMI-1640 medium (Corning, #45000-396) containing 20% FBS (Omega Scientific, #FB-02) and 0.5% penicillin-streptomycin sulfate (Gibco, 10378016). Splenocytes were incubated with 100 U/ml IL-2 (BioLegend, #575404) for 24hr, followed by 300 ng/mL OVA peptide for 24Lhr. Then CD8^+^ T Cells were isolated using the MojoSort™ Mouse CD8^+^ T Cell Isolation Kit (Biolegend, #480035), counted, and resuspended in RPMI-1640 20% FBS medium. Media from MOC2 cells was removed and replaced with 100 μL of fresh culture media without antibiotics, followed by 100 μL of CD8^+^ T cells in media. The co-cultured cells were incubated for 48Lhr. The media and suspended CD8+ T cells were removed and MOC2 cell viability was measured by MTT assay as described above.

### Bulk RNA-Seq data analysis and GSEA

Raw RNA-Seq reads were aligned to the human genome using HISAT2. Gene-level read quantification was performed with htseq-count using GENCODE annotations^45,46^. FPKM values were calculated for normalization, and differential gene expression analysis was conducted using the DESeq2 R package^47^.

GSEAPreranked was performed using the fold change as the input and either the cancer hallmark gene sets or hEMT-related gene set as the library^48^. The enriched terms with FDRL<L0.05 were identified.

### In silico compound sensitivity screen

To identify compounds selectively targeting ETS1-high cancers, we conducted an in silico correlation analysis using the PRISM Primary Repurposing dataset^24^, which reports sensitivity profiles of 4,687 compounds tested across more than 483 human cancer cell lines. Matched RNA-Seq data were obtained from CCLE data^22^ to quantify ETS1 expression in each cell line. For each compound, Pearson correlation coefficients were calculated between ETS1 expression levels and the area under the dose–response curve (AUC) values, where a negative correlation indicated higher sensitivity in ETS1-high lines. Data processing and visualization were performed in R (v4.2.2) using tidyverse and ggplot2 packages.

### ChIP-Seq data analysis

Raw ChIP-Seq reads were mapped to the human genome using bwa, followed by conversion to sorted BAM files with SAMtools^49,50^. PCR duplicates were removed using sambamba, and blacklist regions were excluded with BEDtools^51–53^. Peaks were called using MACS2; for H3K27ac ChIP-seq, the parameter “-nomodel --extsize 146” was utilized, while default parameters were applied to ETS1 ChIP-seq with default parameters^54^. BigWig files were generated with bamCompare (CPM normalization) from deepTools^55^.

### Intratumoral heterogeneity (ITH) analysis using single-cell RNA-Seq (scRNA-seq) data

ITH analysis and identification was performed using scRNA-seq data as described recently^9,56^. Briefly, non-negative matrix factorization (NMF) was applied to each of 16 UASCC cell lines with matched Metmap and scRNA-seq data. Expression matrices were adjusted to exclude negative values, and NMF was performed with factors (K) ranging from 6 to 9, generating 30 gene modules per cell line. Gene modules were filtered and refined based on overlap criteria within and across cell lines to ensure robustness and reduce redundancy. Ultimately, 50 consensus gene modules were identified, clustered into 7 clusters ITH programs using Jaccard similarity, and signature genes were defined as those present in at least 25% of constituent programs.

To interpret these 7 ITH programs, we compared them to 41 curated meta-programs (MPs) derived from scRNA-seq of pan-cancer tumors^6^. Similarity was evaluated using the Jaccard index and single-cell score correlation for each ITH program against the MPs. Statistical significance was determined by permuting the ITH programs 100 times, with a confidence threshold set at 99.9%.

To score the ITH programs in each single cell, we generated 1,000 background gene sets with comparable expression levels for each consensus program, as described previously^6^. For each cell, we calculated the average centered expression of the ITH program and background sets. The p-value was defined as the proportion of background sets with higher expression than the ITH program. The ITH program score was then calculated as −log10(p) and linearly rescaled to [0,1].

### Data availability

RNA-Seq data generated in this study have been deposited into GEO repository under accession number GSE300573.

