## Supplemental Figures for "ETS1 Orchestrates a Hybrid EMT Program Driving in vivo Metastasis and Immune Evasion"

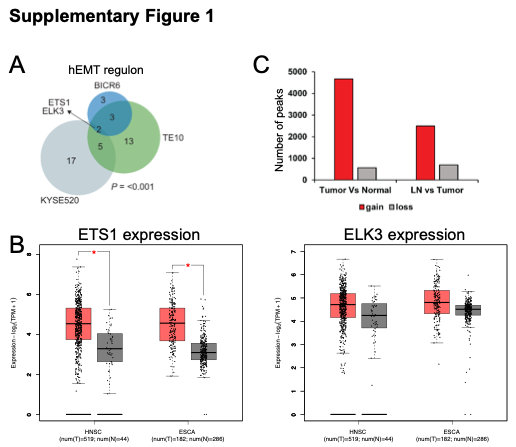

**Supplementary Figure 1.** (A) A Venn diagram showing SCENIC hEMT regulon predictions in three UASCC cell lines. (B) The mRNA expression levels of ETS1 and ELK3 from TCGA cohorts, analyzed via gepia2. Red bar represents tumor samples and grey bar represents normal samples. (C) Differential H3K27ac peak analysis between normal samples, primary ESCC and LN metastases, corresponding to Figure 2O.

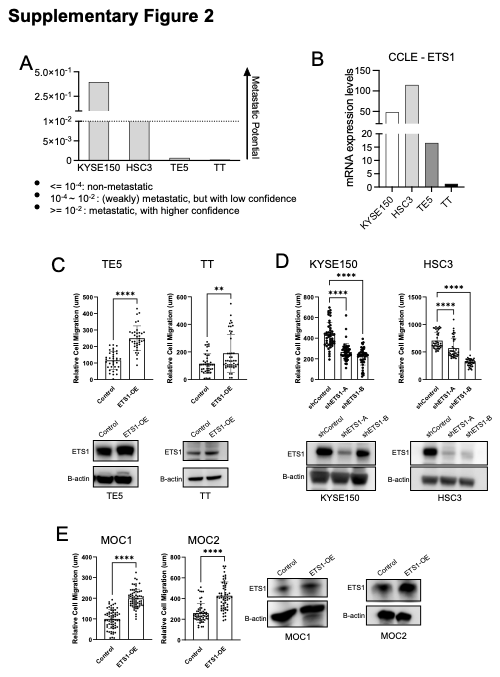

**Supplementary Figure 2.** (A) Column plots showing the mean values of metastatic potential of indicated cell lines. (B) Column plots showing the expression of ETS1 mRNA of indicated cell lines. (C-E) Bar graphs show the quantification of relative cell migration 24 hours post wound and represent the mean ± SD of three experimental replicates. **P<0.01, ****P<0.0001; P-values were determined using an unpaired t-test (or one-way ANOVA with multiple comparisons. Western Blotting validating ETS1 changes in the corresponding experiments.

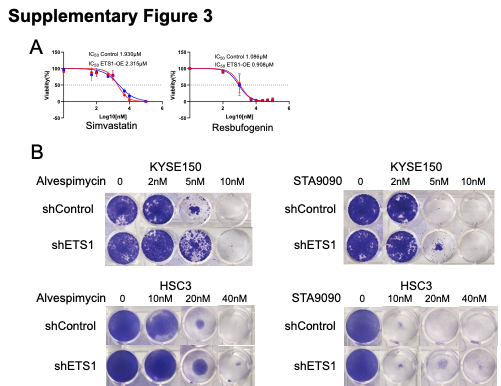

**Supplementary Figure 3.** (A) Cell viability (MTT) IC50 assays in both control (red) and ETS1 overexpressing (blue) UASCC cell lines. Compounds are represented in nM using the Log10 scale. Data points represent the mean ± SD of at least two experimental replicates. (B) Colony formation assays upon exposure to Alvespimycin and STA9090 in control or ETS1-knockdown UASCC cell lines.

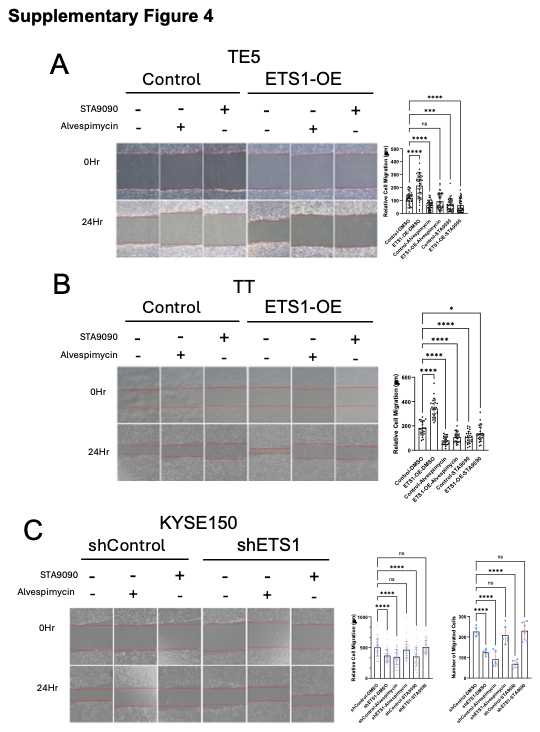

**Supplementary Figure 4.** (A-C) Wound healing assays in control, ETS1-overexpressing or ETS1-knockdown UASCC cell lines with or without the exposure to Alvespimycin or STA9090. Bar graphs show the quantification of relative cell migration and represent the mean ± SD of three experimental replicates. *P<0.05, ***P<0.001, ****P<0.0001; P-values were determined using a one-way ANOVA with multiple comparisons.

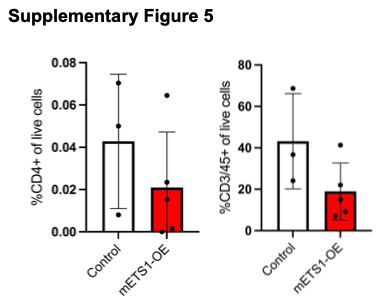

**Supplementary Figure 5.** The proportion of CD3+ and CD4+ T cells out of total live cells from draining lymph nodes, measured by flow cytometry analysis. Bars represent mean ± SD of independent samples.
